# Effects of mutation and selection on plasticity of *TDH3* promoter activity in *Saccharomyces cerevisiae*

**DOI:** 10.1101/173344

**Authors:** Fabien Duveau, David C. Yuan, Brian P.H. Metzger, Andrea Hodgins-Davis, Patricia J. Wittkopp

## Abstract

Phenotypic plasticity is an evolvable property of biological systems that can arise from environment-specific regulation of gene expression. To better understand the evolutionary and molecular mechanisms that give rise to plasticity in gene expression, we quantified the effects of 235 single nucleotide mutations in the *Saccharomyces cerevisiae TDH3* promoter (*P_TDH3_*) on the activity of this promoter in media containing glucose, galactose, or glycerol as a carbon source. We found that the distributions of mutational effects differed among environments because many mutations altered the plastic response exhibited by the wild type allele. Comparing the effects of these mutations to the effects of 30 *P_TDH3_* polymorphisms on expression plasticity in the same environments provided evidence of natural selection acting to prevent the plastic response in *P_TDH3_* activity between glucose and galactose from becoming larger. The largest changes in expression plasticity were observed between fermentable (glucose or galactose) and nonfermentable (glycerol) carbon sources and were caused by mutations located in the RAP1 and GCR1 transcription factor binding sites. Mutations altered expression plasticity most frequently between the two fermentable environments, however, with mutations causing significant changes in plasticity between glucose and galactose distributed throughout the promoter, suggesting they might affect chromatin structure. Taken together, these results provide insight into the molecular mechanisms underlying gene-by-environment interactions affecting gene expression as well as the evolutionary dynamics affecting natural variation in plasticity of gene expression.

**Significance Statement:** From seasonal variation in the color of butterfly wings to trees bending toward the light, organisms often change in response to their environment. These changes, known as phenotypic plasticity, can result from differences in how genes are expressed among environments. Mutations causing environment-specific changes in gene expression provide raw material for phenotypic plasticity, but their frequency, effect size and direction of effects among environments are not well understood. This study shows that mutations in the promoter of a yeast metabolic gene often display environment-dependent effects on gene expression and that these environment-dependent effects have been shaped by selection in natural populations.

## Introduction

Phenotypic plasticity, which is the ability of an organism to develop different phenotypes in different environments, has been observed for diverse traits in diverse species [1, 2]. This plasticity can facilitate adaptation to dynamically changing environments [3] but is not always adaptive [4, 5]. Indeed, the role of natural selection in the evolution of phenotypic plasticity has long been the subject of debate (reviewed in [6]). For selection to act on plasticity, populations must exhibit genetic variation responsible for differences in plasticity among individuals [7]. Such genetic variation, frequently detected as significant gene-by-environment interactions (GxE) in studies of quantitative traits [8], results from the mutational process generating variation in phenotypic plasticity, natural selection changing frequencies of alleles based on their effects on fitness, and genetic drift changing allele frequencies stochastically based on population size. Understanding how new mutations generate variation in plasticity and how selection acts on this variation are thus key for understanding the origin, maintenance, and evolution of phenotypic plasticity [9-12].

One common source of phenotypic plasticity is environment-specific regulation of gene expression (reviewed in [13]), which has been described for many organisms in response to different environmental cues (*e.g.*, [14-18]). Standing genetic variation altering these environment-specific responses also appears to be common (*e.g*., [18-22]). For example, a study comparing gene expression in two strains of the baker’s yeast *Saccharomyces cerevisiae* grown in media containing glucose or ethanol as a carbon source found that 79% of genes showed expression differences attributed to the environment and 47% of genes had expression affected by quantitative trait loci (QTL) with evidence of significant interactions with the environment [20]. QTL that alter plasticity of gene expression (and therefore contribute to GxE) appear to be common in other systems as well [18, 22, 23]. In most cases, it remains unclear whether the variation in gene expression plasticity observed has resulted from the neutral accumulation of mutations or has been shaped by the action of natural selection.

To separate the contributions of mutation and selection, mutation accumulation experiments have been used to isolate the effects of spontaneous mutations on a variety of quantitative traits (reviewed in [11, 12, 24]), including gene expression (*e.g*., [25-28]). By comparing trait variation among lines in which non-lethal mutations have accumulated in the near absence of selection, these studies estimate the mutational variance (V_m_), which is the phenotypic variance added to a population each generation by new mutations. Mutational variance was found to differ among environments (indicating GxE) for some traits [29-32] but not others [33, 34], demonstrating that the propensity of new mutations to alter plasticity differs depending on the trait and environments considered. Because mutation accumulation experiments allow variation to accumulate throughout the genome, trait variation observed in these studies cannot easily be tied to a particular mutation(s) [28, 35, 36]. Recent studies have provided more direct links between specific mutations and variation in gene expression by using random or targeted mutagenesis to mutate the *cis*-regulatory region (promoter or enhancer) of a focal gene and then using high-throughput RNA sequencing or fluorescence reporters to determine the effects of these mutations on expression [37-41]. By empirically describing the effects of new mutations on gene expression, a neutral model is generated that can be compared to the effects of genetic variants segregating in natural populations to infer the effects of natural selection [26, 42-44]. To date, all such comparisons have been made between the effects of mutations and polymorphisms in a single environment; however, a similar approach can be used to disentangle the relative contributions of mutation and selection to variation in gene expression plasticity if the effects of mutations and polymorphisms are measured in multiple environments.

Here, we use this approach to examine the effects of mutation and selection on plasticity of a *cis*-regulatory sequence controlling gene expression by measuring the effects of 235 mutations and 30 polymorphisms in the *S. cerevisiae TDH3* promoter in media containing one of three different carbon sources (glucose, galactose, and glycerol). This promoter comes from the *TDH3* gene, which encodes a glycolytic enzyme (glyceraldehyde-3-phosphate dehydrogenase) involved in metabolism of fermentable sugars such as glucose and galactose as well as non-fermentable carbon sources such as glycerol (Figure 1A). These three carbon sources are likely to be encountered and used by natural populations of *S. cerevisiae* (albeit at unknown frequencies), since the genome contains conserved sets of genes for metabolizing glucose [45], galactose [46], and glycerol [47]. Genes required specifically for metabolism of galactose or glycerol are repressed in presence of glucose, which appears to be the preferred carbon source for *S. cerevisiae* [48]. We found that activity of the wild type *TDH3* promoter displays plasticity among environments containing glucose, galactose or glycerol as a carbon source. We also found that the distributions of effects of *cis*-regulatory mutations on promoter activity varied among these three environments because of differences in the frequency, magnitude and direction of GxE effects observed between pairs of environments. When we compared the effects of these mutations to the effects of *P_TDH3_* polymorphisms segregating in natural populations, we found that mutations tended to cause larger differences in promoter activity between the two fermentable environments than polymorphisms, suggesting that natural selection has preferentially eliminated genetic variants that increase expression plasticity between glucose and galactose. Finally, by comparing the locations of mutations that showed environment-dependent effects on expression to the locations of previously identified functional elements in the *TDH3* promoter, we found evidence suggesting that plasticity in the activity of this promoter is mediated by different molecular mechanisms in different environments.

**Figure 1.**
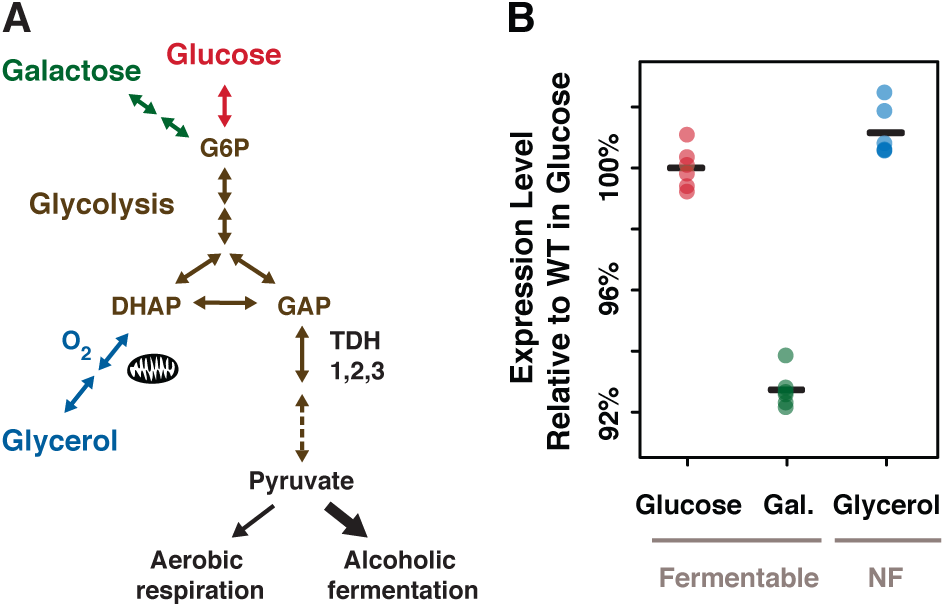
TDH3 functions in central metabolism and its promoter activity is plastic among environments. (A)*TDH3* encodes one of *S. cerevisiae’s* three glyceraldehyde-3-phosphate dehydrogenase (GAPDH) proteins involved in glycolysis and required for metabolism of fermentable carbon sources such as glucose and galactose and of the non-fermentable carbon source glycerol after aerobic conversion in dihydroxyacetone phosphate (DHAP) in mitochondria. For greater detail, see (http://pathway.yeastgenome.org/; [46, 47, 83]). (B) Activity of the wild type *TDH3* promoter in media containing glucose (red), galactose (green), and glycerol (blue) is shown. Colored dots represent the median fluorescence level of each of the six replicate populations, and the black bars indicate the mean of the six median fluorescent levels observed for each environment.

## Results and Discussion

### *Plasticity of* TDH3 *promoter activity*

To measure *TDH3* promoter (*P_TDH3_*) activity in living cells, we quantified the expression of a *P_TDH3_*-*YFP* reporter gene inserted in the genome of the laboratory strain YPW1 that contains a wild type allele of *P_TDH3_* fused to the coding sequence of the yellow fluorescent protein (YFP Venus) [49]. The fluorescence level of YPW1 was measured after growth in rich media containing glucose, galactose, or glycerol as a carbon source to quantify differences in promoter activity among environments and measure plasticity. For each environment, the level of fluorescence of at least 10,000 cells was quantified by flow cytometry in each of six replicate populations, and the median fluorescence of each sample was divided by the average fluorescence measured among the six replicates in glucose (“Plasticity.WT” worksheet in Dataset S1). We observed statistically significant differences in expression of the reporter gene (*i.e.*, expression plasticity) between each pair of environments (Figure 1B; t-tests, P_glu-gal_ = 3.2 × 10^−9^, P_glu-gly_ = 0.025, P_gal-gly_ = 6.1 × 10^−9^). Surprisingly, a larger difference in expression level was observed between the two fermentable carbon sources, glucose and galactose (7.3% lower expression in galactose), than between one of the fermentable carbon sources (glucose) and the non-fermentable (glycerol) carbon source (1.2% higher expression in glycerol), despite the fact that growth in the two fermentable carbon sources involves more similar metabolic processes than growth in fermentable and non-fermentable environments (Figure 1A).

### Environment-dependent distributions of mutational effects

To determine how new mutations altered the plastic response observed for the wild type allele of the *TDH3* promoter, we examined 235 mutant versions of the reporter gene created by using site-directed mutagenesis to change one of the 241 Gs and Cs in *P_TDH3_* into an A or T, respectively, in each strain [42] (Figure 2A). Each of these 235 mutant genotypes, as well as the un-mutated “wild type” genotype, was then grown in six replicate populations in media containing glucose, galactose, or glycerol as a carbon source (Figure 2B,C). The activity of the *P_TDH3_*-*YFP* reporter gene was assayed in ~8000 cells on average in each population using flow cytometry (Figure 2D). After normalizing for differences in cell size and controlling for technical variation, the median fluorescence level was calculated for each population and used as a proxy for promoter activity (Figure 2E; “All.Mutations.Data” worksheet in Dataset S1).

**Figure 2.**
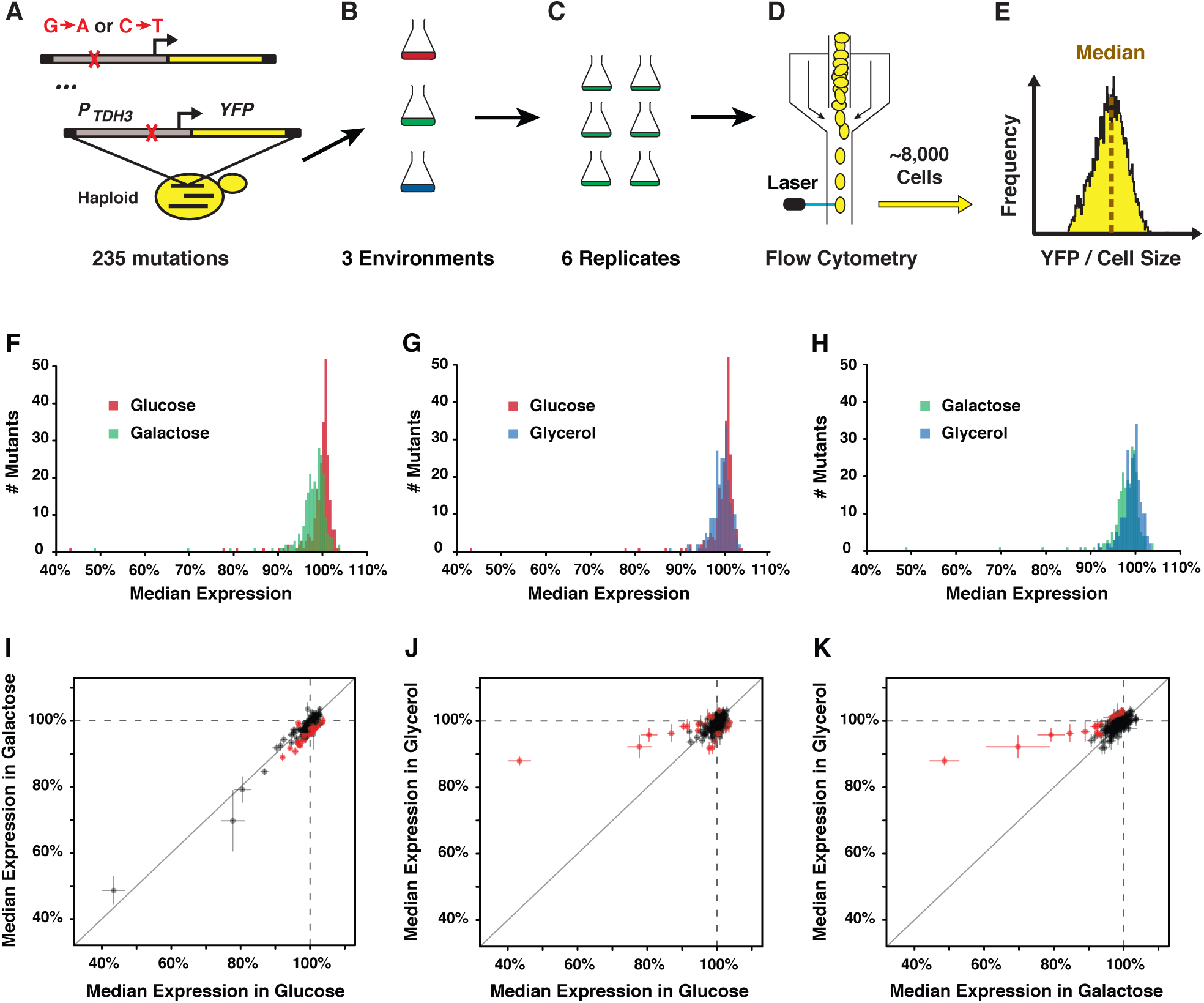
Effects of 235 *cis*-regulatory mutations on activity of the *TDH3* promoter in glucose, galactose, and glycerol. (A-E) Experimental overview. (A) Site-directed mutagenesis was used to change Gs to As and Cs to Ts in the *TDH3* promoter (*P_TDH3_*) driving expression of a yellow fluorescent protein (YFP) in each of 235 strains [42]. (B) Each of these strains, plus a strain carrying the wild type *P_TDH3_*-*YFP* allele, was then grown in three environments containing different carbon sources. (C) Six replicate populations for each genotype were analyzed in each type of media. (D) Fluorescence of individual cells in each population was assessed using flow cytometry. (E) After correcting for differences in cell size, the median fluorescence was determined for each population. (F-H) Histograms showing the median activity (averaged across six replicate populations) for the 235 mutant *TDH3* promoters relative to the average activity of the wild type promoter in the same environment for cells grown in (F) glucose (red) or galactose (green), (G) glucose (red) or glycerol (blue) and (H) galactose (green) or glycerol (blue). (I-K) Comparisons of effects of *TDH3* cis-regulatory mutations between (I) glucose and galactose, (J) glucose and glycerol and (K) galactose and glycerol. The median activity of mutant promoters is expressed relative to the average activity of the wild type allele in each environment, with dots showing the average activity across six replicates and error bars showing 95% confidence intervals. Dots colored red showed evidence of a statistically significant gene-by-environment interaction based on t-tests corrected for multiple testing using the Benjamini-Hochberg False Discovery Rate adjustment (P_adj_ < 0.01).

To quantify changes in plasticity caused by mutations, we controlled for the plasticity of the wild type *P_TDH3_* allele (Figure 1B) by dividing the median fluorescence level of each replicate population of the 235 mutant genotypes by the average median fluorescence measured for the wild type allele in the corresponding environment. The mean relative fluorescence from six replicates was then used to estimate the effect of each genotype on promoter activity in each environment, with variability among replicates used to calculate the 95% confidence interval around this mean. Using this relative measure of expression, we found that the distributions of effects of the 235 mutant genotypes on *P_TDH3_*-*YFP* expression were different between glucose and galactose (Figure 2F; KS test with permutations, D = 0.43, P < 5 × 10^−6^), between glucose and glycerol (Figure 2G; KS test with permutations, D = 0.23, P < 5 × 10^−6^) and between galactose and glycerol (Figure 2H; KS test with permutations, D = 0.22, P < 5 × 10^−6^). In galactose, the distribution of mutational effects was more skewed toward decreases in expression (median effect = −1.5%) than in glucose (median effect = +0.3%, permutation test, P_glu-gal_ < 5 × 10^−6^) or glycerol (median effect = −0.5%; permutation test, P_gly-gal_ < 5 × 10^−6^). This mutational bias toward lower expression suggests that new mutations in the *TDH3* promoter will tend to further decrease *TDH3* expression in galactose relative to its expression in glucose or glycerol. As a consequence, newly arising variation in this promoter sequence is predicted to exacerbate the plasticity observed for the wild type allele.

In addition to differences in their median effects, the distributions of mutational effects also displayed different degrees of dispersion depending on the environment (Figure 2F-H). To quantify the dispersion of each distribution, we calculated the median absolute deviation (MAD) of the median expression levels observed across the 235 mutant strains in each environment. The MAD is a measure of dispersion less sensitive to rare outliers than the standard deviation. We found that the variability of mutational effects was similar in galactose and glycerol (Figure 2H; MAD_gal_ = 1.97%, MAD_gly_ = 1.75%; permutation test, P_gal-gly_ = 0.12), but lower in glucose (MAD_glu_ = 1.12%) than in galactose (Figure 2F; permutation test, P_glu-gal_ < 5 × 10^−6^) or glycerol (Figure 2G; permutation test, P_giu-giy_ = 3.3 × 10^−4^). In addition, the median effect size of mutations on promoter activity was lowest in glucose (median absolute effect of 235 mutations: glucose = 0.92%; galactose = 1.80%; glycerol = 1.16%; permutation tests, P_glu-gal_ < 5 × 10^−6^, P_glu-gly_ = 0.036, P_gal-gly_ = 3 × 10^−5^), indicating that mutant promoters conferred expression levels closest to the wild type promoter in glucose. Taken together, these observations suggest that the activity of the *TDH3* promoter is more robust to the effects of new *cis*-regulatory mutations in glucose, which is the preferred carbon source of *S. cerevisiae* [48], than in glycerol or galactose.

### Frequency, magnitude and direction of gene-by-environment interactions

Differences in the distributions of mutational effects among the three environments indicated that at least some of these mutations in the *P_TDH3_* promoter have environment-dependent effects. To identify these mutations (*i.e.*, to test for gene-by-environment interactions), we compared the effects of each mutation relative to the wild type allele between environments using a series of pairwise t-tests with a Benjamini-Hochberg False Discovery Rate (FDR) correction for multiple testing. We found that the effects of individual mutations were more strongly correlated between the two fermentable carbon sources (Figure 2I, Pearson correlation coefficient: r_glu-gal_ = 0.94) than between either fermentable carbon source and the non-fermentable glycerol (Figure 2J, r_glu-gly_ = 0.55; Figure 2K, r_gal-gly_ = 0.65). However, nearly three times as many mutations showed statistically significant evidence of GxE for the comparison between glucose and galactose than for other pairs of environments (P_env1-env2_ < 0.01: N_glu-gal_ = 75, N_glu-gly_ = 29, N_gal-gly_ = 26). This disconnect results from differences in the magnitude of significant GxE effects between pairs of environments, with the largest differences observed when comparing fermentable and nonfermentable environments (Δ_glu-gly_ = 6.3%, Δ_gal-gly_ = 6.0%, Δ_glu-gal_ = 3.1%; Mann-Whitney-Wilcoxon tests P_Δglu-gly *vs* Δgal-gly_ =0.77; P_Δglu-gly *vs* Δglu-gal_ = 0.0032; P_Δgal-gly *vs* Δglu-gal_ = 0.029). Interestingly, these data indicate that mutations with large GxE effects can arise when plasticity of the wild type allele is small (glucose *vs* glycerol) and mutational effects can be well correlated between environments even when the plasticity of the wild type allele is large (glucose *vs* galactose). Biases in the direction of GxE effects also differed among pairwise comparisons: 73 of 75 mutations with significant GxE effects between glucose and galactose (Figure 2I) and all 24 mutations with significant GxE effects between glycerol and galactose (Figure 2K) caused lower expression in galactose (binomial test, P < 10^−6^ in both cases), whereas no bias in the direction of GxE effects was observed between glucose and glycerol (Figure 2J; N_glu>gly_ = 15 and N_gly>glu_ = 14; binomial test, P = 1). The strong directional bias of the GxE effects observed between galactose and other carbon sources suggests that plasticity of expression could increase in the absence of natural selection simply through the random occurrence of *cis*-regulatory mutations.

### Effects of selection on expression plasticity

Genetic variation for plasticity segregating in natural populations can be influenced both by the mutational process creating new genetic variation and by natural selection filtering genetic variants based on their phenotypic effects. To test for evidence of selection acting on plasticity of *TDH3* promoter activity, we compared the effects of mutations in media containing glucose, galactose, and glycerol to the effects in the same environments of 30 polymorphisms in this promoter that were identified in 85 strains of *S. cerevisiae.* To quantify the effects of these polymorphisms on promoter activity, 39 haplotypes of *P_TDH3_* that differed from each other by one to ten polymorphisms (Dataset S2) were cloned upstream of the *YFP* coding sequence [42] and the fluorescence levels of the resulting strains were measured by flow cytometry after growth of six replicate populations in each of the three carbon source environments. The effect of each of 30 unique polymorphisms (Figure S1, “Polymorphisms.Effects” worksheet in Dataset S1) was inferred by dividing the fluorescence measured for a haplotype containing that polymorphism by the average fluorescence measured for an ancestral haplotype that differed by only that polymorphism, as described more fully in Metzger et al. [42], Duveau et al. [50] and the Methods section. As a group, these 30 polymorphisms caused expression to vary from 94.6% to 102.5% of the ancestral allele in glucose (Figure 3A), 96.1% to 105.5% in galactose (Figure 3B), and 95.4% to 103.7% in glycerol (Figure 3C). The ten polymorphic G:C -> A:T transitions, thirteen other types of single nucleotide polymorphisms (SNPs) and seven indel polymorphisms showed similar median effects on expression level in all three environments (Figure S1B-D). Despite the narrow range of effects covered by these *P_TDH3_* polymorphisms, their effects were well correlated between environments (Figure S2, Pearson correlation coefficients: r_glu-gal_ = 0.58, P_glu-gal_ = 8.4 × 10^−4^; r_glu-gly_ = 0.42, P_glu-gly_ = 0.022; r_gal-gly_ = 0.72, Pgly = 6.9 × 10^−6^). Evidence of significant GxE effects was observed for 4 of the 30 polymorphisms between glucose and galactose, 2 between glucose and glycerol, and 2 between galactose and glycerol (Figure S2).

**Figure 3.**
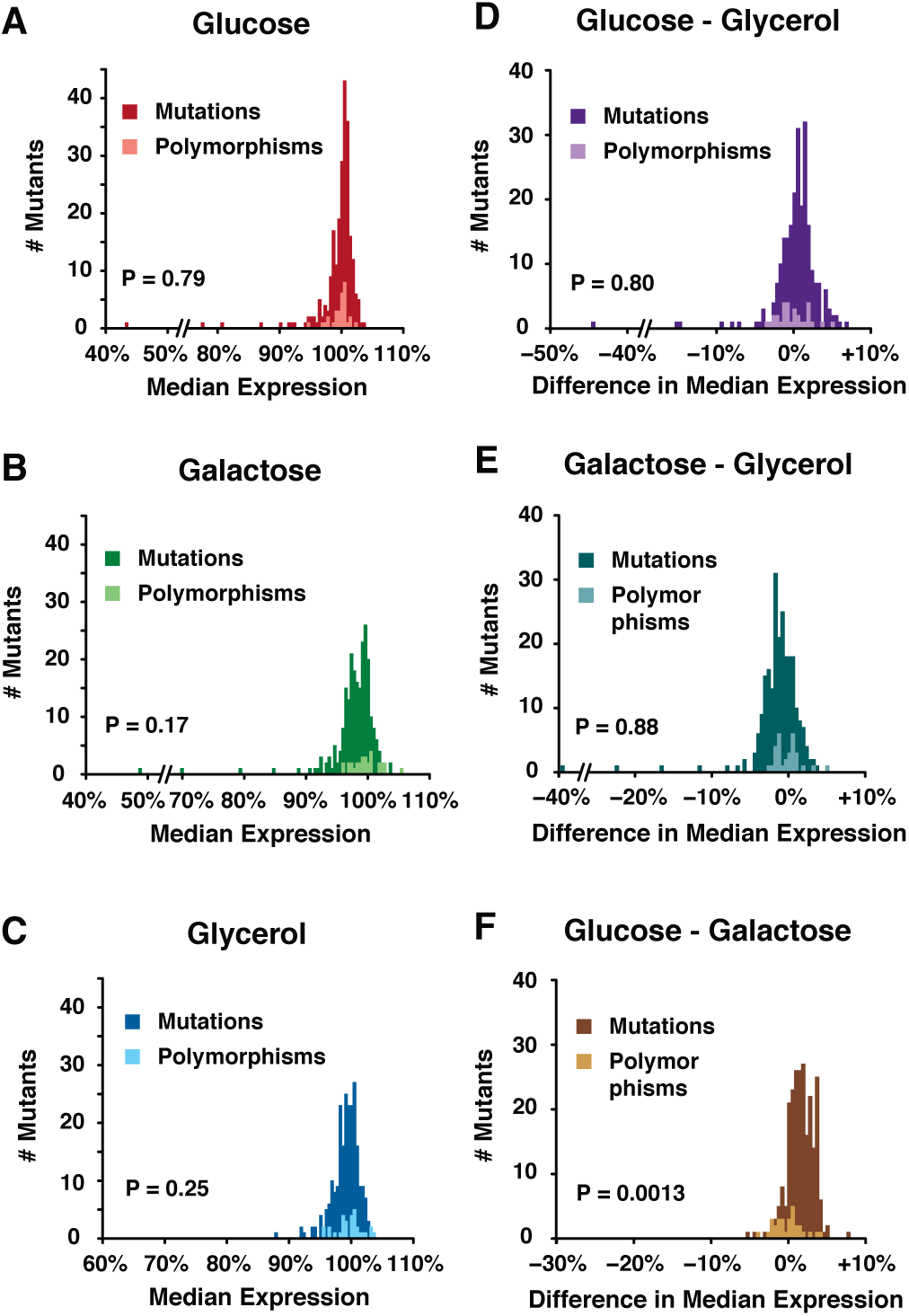
Selection affects natural variation in plasticity of *TDH3* promoter activity. (A-C) Histograms showing the distributions of effects for 235 point mutations and for 30 polymorphisms in the *TDH3* promoter upon growth in (A) glucose, (B) galactose and (C) glycerol. P-values represent the significance of the difference between the effects of polymorphisms and mutations obtained using non-parametric resampling tests as described in the Methods. (D-F) Histograms showing the distributions of differences in effects between environments for 235 point mutations and for 30 polymorphisms in the *TDH3* promoter. The differences of promoter activity were calculated between (D) glucose and glycerol, (E) galactose and glycerol and (F) glucose and galactose. P-values were determined using non-parametric resampling tests to compare the effects of mutations and polymorphisms on plasticity between pairs of environments.

To determine whether selection has maintained genetic variants with a particular subset of effects on expression in natural populations, we compared the distributions of effects of mutations and polymorphisms in each environment with the same non-parametric approach used in Metzger et al. [42]. Briefly, a probability density function was fitted to the empirical distribution of mutational effects in each environment, 100,000 sets of 30 mutations were drawn randomly from each of these functions and the log-likelihood of each set was computed using the corresponding probability density function. The log-likelihoods of the effects observed for the 30 polymorphisms were also determined using the same probability density functions. We then calculated a P-value for the difference in effects between mutations and polymorphisms in each environment by determining the proportion of random sets of mutations with log-likelihood values more extreme than the log-likelihood value calculated for the set of 30 polymorphisms. For cells grown in glucose, we found that the distribution of effects of polymorphisms was not significantly different from random sampling of the mutational distribution (Figure 3A; Figure S3A, P = 0.43), consistent with Metzger et al. [42]. Likewise, we found that the effects of these polymorphisms in galactose (Figure 3B; Figure S3B; P = 0.35) and glycerol (Figure 3C; Figure S3C; P = 0.51) were consistent with random sampling of mutations. Therefore, no significant signature of selection acting on the level of activity of the *TDH3* promoter was detected in any of the three environments tested.

Next, we tested for evidence of selection acting on expression plasticity between environments by comparing the difference of effects between each pair of environments for mutations and polymorphisms using the approach described above. We detected no significant difference in the plasticity of expression conferred by the set of polymorphisms and random sets of mutations between fermentable and non-fermentable environments (glucose and glycerol: Figure 3D; Figure S3D; P = 0.80; galactose and glycerol: Figure 3E; Figure S3E; P = 0.88); however, we observed a significant difference in plasticity between the two fermentable environments, glucose and galactose (Figure 3F; Figure S3F; P = 0.0013), with the 30 polymorphisms showing smaller median differences in expression between glucose and galactose than random sets of 30 mutations (Δpol_(glu.gal)_ = +0.08% *vs* Δmut_(glu.gal)_ = +1.64%; permutation test, P = 5.2 × 10^−3^). This observation suggests that selection has preferentially eliminated new mutations that cause the largest expression plasticity between glucose and galactose. Selection might have also disfavored mutations with a large impact on plasticity observed between fermentable and non-fermentable environments (Figure 2J,K), but the low frequency of these mutations suggests that a larger number of polymorphisms would be needed to detect selection acting on plasticity between fermentable and non-fermentable environments.

Observing evidence of selection acting on plasticity between glucose and galactose without also observing evidence of selection acting on expression levels in either environment individually was surprising. To better understand this result, we examined the effects of mutations and polymorphisms in glucose and galactose more closely. We found that mutations tended to confer slightly higher expression levels than polymorphisms in glucose (difference of median effects: Δglu_(mut-pol)_ = +0.37%) and lower expression levels than polymorphisms in galactose (difference of median effects: Δgal_(mut-pol)_ = −1.0%). Because these (weak) directional biases were in opposite directions and because polymorphisms had similar effects in glucose and galactose (difference of median effects: Δpol_(glu-gal)_ = +0.08%; permutation test, P = 0.85), the difference in effects between mutations and polymorphisms was larger and easier to detect between the two environments than in either environment alone. These observations suggest that selection might disfavor mutations that increase expression in glucose and/or decrease expression in galactose. The larger difference in median effects between mutations and polymorphisms in galactose than glucose as well as the minimal fitness effects of small increases in *TDH3* expression in glucose reported recently [50] suggest that selection against mutations that decrease expression in galactose is more likely to be driving this pattern.

### Potential molecular mechanisms underlying gene-by-environment interactions

One of the reasons that we selected the *TDH3* promoter for this work was that prior studies had characterized functional elements within its sequence (Figure 4A), providing an opportunity to develop hypotheses about the molecular mechanisms underlying plasticity and gene-by-environment interactions. Specifically, Kuroda et al. [51] described an upstream activating sequence (UAS1, 426 to 528 bp upstream of the start codon) as required for expression in a fermentable carbon source (glucose) but not a non-fermentable carbon source (glycerol + lactate). This sequence contains binding sites for the RAP1 and GCR1 transcription factors [52-54]. An upstream repressing sequence (URS, 419 to 431 bp upstream of the start codon) adjacent to UAS1 was also identified by Kuroda et al. [51] that appeared to repress activity of a second upstream activating sequence (UAS2) in the fermentable environment assayed (Figure 4A). This UAS2 (255 to 309 bp upstream of the start codon) was described as being primarily responsible for activation in the non-fermentable environment tested [51].

**Figure 4.**
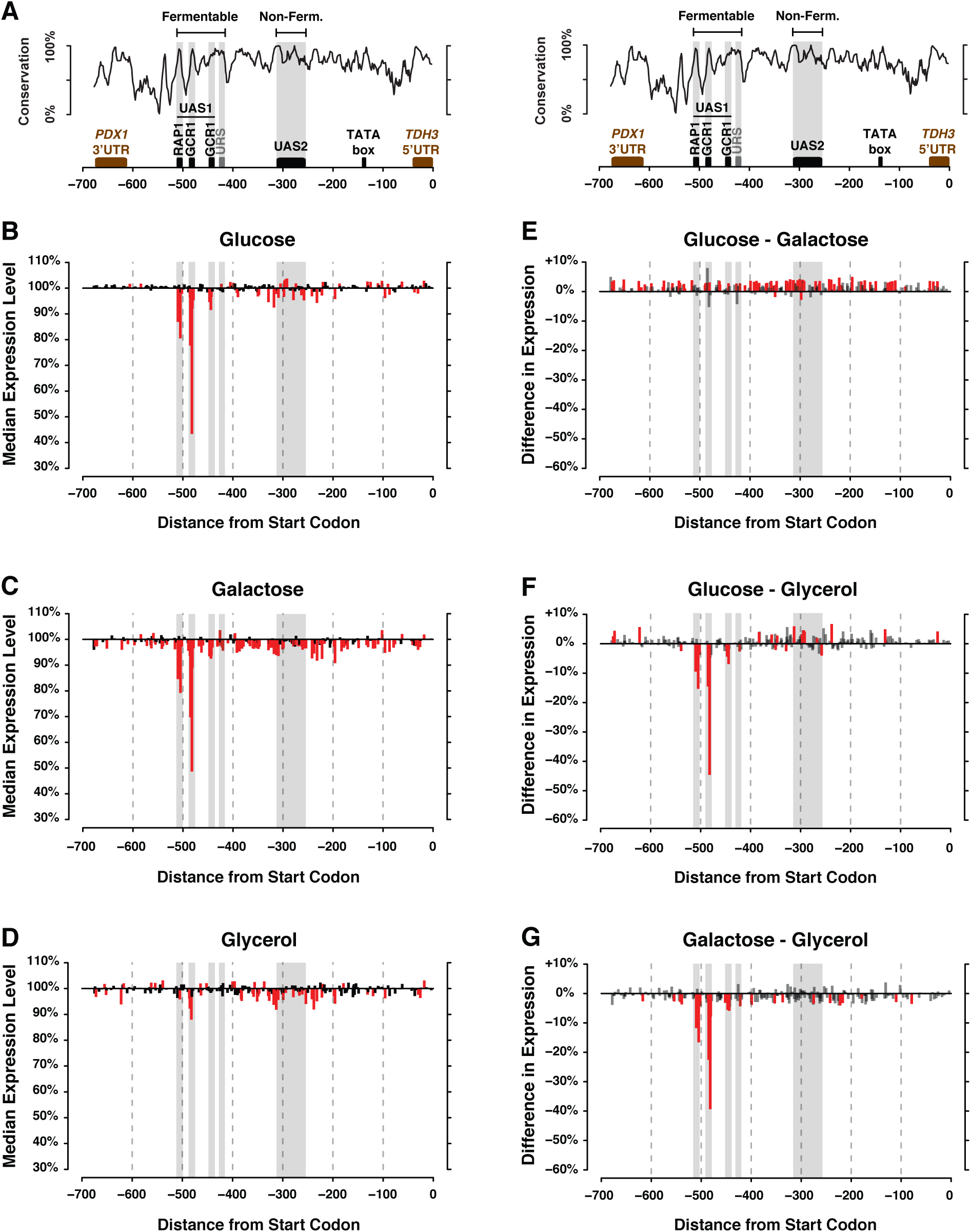
Locations of mutations affecting activity and plasticity of the *TDH3* promoter within the promoter sequence. (A) A summary of previously identified functional elements in the *TDH3* promoter is shown. These elements include binding sites for the RAP1 and GCR1 transcription factors as well as sequences shown previously by deletion analysis to be required for expression during growth on a fermentable carbon source, glucose, (UAS1 and URS) or a non-fermentable carbon source, glycerol plus lactate (UAS2) [51]. For orientation, the TATA box is also indicated, although this sequence was not mutated in any of the strains analyzed. The black curve shows sequence conservation across species of the *Saccharomyces senso stricto* genus. (B-D) The effect of each mutation on the median expression level of *P_TDH3_*-*YFP* relative to expression of the unmutated wild type allele is shown for cells grown on (B) glucose, (C) galactose, and (D) glycerol. Mutation effects represented in red led to a significant change in expression relative to the wild type allele (t-test, P < 0.01). (E-G) The difference in effect of each mutation on *P_TDH3_*-*YFP* in each pair of environments relative to the wild type allele is shown for (E) glucose and galactose, (F) glucose and glycerol, and (G) galactose and glycerol. Mutation represented in red showed significant difference of effects between the two environments based on t-tests corrected for multiple testing using the Benjamini-Hochberg False Discovery Rate adjustment (P_adj_ < 0.01). Areas shaded in grey correspond to functional elements located directly above these regions in panel A.

To better understand the environment-specific regulation of the *TDH3* promoter, we compared the locations of mutations with significant effects on *P_TDH3_* activity to these previously described functional elements. In the two fermentable environments we examined (glucose and galactose), the largest effect mutations altered sequences in the RAP1 and GCR1 transcription factor binding sites (Figure 4B, C). The effects of mutations in these binding sites were much smaller in the non-fermentable environment (glycerol), although they remained among the sites with the largest effects on expression even in this environment (Figure 4D). This result is consistent with the UAS1 region playing a larger role in regulating *TDH3* expression in fermentable than nonfermentable carbon sources, but also suggests that activity of UAS1 is not strictly limited to fermentable environments. In glycerol, many mutations that affected activity of the *TDH3* promoter were concentrated near the UAS2 region previously described as required for expression in media containing glycerol and lactate [51] but mutations with significant effects also extended beyond this region (Figure 4D), indicating that the functional region used for expression in glycerol is larger than the UAS2 region described by Kuroda et al. [51]. Mutations in the UAS2 region also impacted expression in glucose and galactose, indicating their function is not restricted to non-fermentable carbon sources. In the repressive URS sequence, one mutation (423 bp upstream of the start codon) increased expression in glucose and another mutation (426 bp upstream of the start codon) increased expression in galactose, but none had significant effects in glycerol (Figure 4B-D), consistent with its previous description as a repressor in fermentable carbon sources. Additional mutations in this sequence had significant effects only in galactose, where they decreased expression (Figure 4B-D).

To identify molecular mechanisms that might give rise to gene-by-environment effects of single nucleotide changes in the *TDH3* promoter, we mapped the differences of mutational effects between each pair of environments on the promoter architecture. In the comparison between the two fermentable environments (glucose and galactose), the 75 mutations with significantly different effects between glucose and galactose (73 of which showed higher expression in glucose than galactose) appeared to be distributed randomly throughout the promoter (Figure 4E). The result of a nearest neighbor statistical test for clustering of mutations was consistent with this observation (standard deviation of distance to nearest neighbor: observed = 6.63 *vs* expected = 8.12; P = 0.51). Between either fermentable carbon source (glucose or galactose) and the non-fermentable glycerol, mutations with the largest GxE effects (expression differences ≥ 2% between environments) appeared to be concentrated in previously identified functional elements (Figure 4F,G). In the comparison between glucose and glycerol (Figure 4F), permutation testing confirmed that the previously identified UAS1 sequence required for expression on fermentable carbon was significantly enriched for mutations showing evidence of gene-by-environment interactions (Figure 4F, number of mutations with significant GxE effects in UAS1: observed = 9 *vs* expected = 4.4; P = 0.009), as was the UAS2 element previously shown to be required for expression during growth on glycerol plus lactate (number of mutations with significant GxE effects in UAS2: observed = 6 *vs* expected = 3.0; P = 0.027). In the comparison between galactose and glycerol (Figure 4G), significant enrichment of mutations showing evidence of gene-by-environment interactions was observed for the UAS1 region (number of mutations with significant GxE effects in UAS1: observed = 9 *vs* expected = 4.1; P=0.001) but not for the UAS2 region (number of significant GxE effects in UAS2: observed = 1 *vs* expected = 3.0; P = 0.94).

Taken together, these data suggest that different molecular mechanisms underlie gene-by-environment interactions in different pairs of environments. Transcription factor binding sites for RAP1 and GCR1 in UAS1, and potentially binding sites for other transcription factors in UAS2, appear to regulate *TDH3* expression differently in fermentable (glucose or galactose) and nonfermentable (glycerol) environments. The fact that mutations with the largest GxE effects between either pair of fermentable and non-fermentable carbon sources affected the RAP1 and GCR1 binding sites (Figure 4F,G) is consistent with this hypothesis. By contrast, mutations with significant GxE interactions between the two fermentable environments (glucose and galactose) were distributed throughout the *TDH3* promoter (Figure 4E), suggesting that they might affect more widespread environment-specific chromatin structure rather than specific binding sites. Pavlovic and Hörz [55] identified a nucleosome-free region in *P_TDH3_* extending from the 5’ end of UAS1 to the 3’ end of UAS2 in both glucose and glycerol but did not examine chromatin structure in galactose. We hypothesize that this region exhibits altered nucleosome positioning when cells are grown in galactose. Differences in nucleosome occupancy between these environments may reduce expression of the wild type allele in galactose relative to glucose (Figure 1B) while maintaining the strong correlation of mutational effects observed between these two environments (Figure 3I). Changes in chromatin structure across *P_TDH3_* in galactose could also explain why so many mutations distributed throughout the promoter reduced expression 2 to 5% only in galactose (Figure 4C). Consistent with this model of environment-specific nucleosome occupancy, prior work has shown that activity of promoters occupied by nucleosomes tends to be affected by mutations distributed throughout the whole promoter sequence whereas activity of promoters with a nucleosome-free region upstream of the transcription start site tends to be affected by a smaller number of mutations clustered in transcription factor binding sites [56]. In addition, differences in nucleosome positioning have been observed for several promoters between yeast cells grown in glucose and galactose [57-59].

## Conclusions

By characterizing the effects of 235 single nucleotide changes in the *TDH3* promoter on its activity in three different environments, we have shown that changes in the environment can modify the distribution of expression phenotypes generated by new cis-regulatory mutations. We observed mutational biases toward decreased *P_TDH3_* activity in galactose and (to a lesser extent) in glycerol, but not in glucose, suggesting that the random accumulation of mutations would tend to increase the plastic response of *P_TDH3_* activity in these environments relative to the wild type allele of the promoter. In addition, the variability of effects observed among the 235 *P_TDH3_* mutations was lower in glucose than in galactose or glycerol, indicating that mutational robustness was greatest in the environment containing the preferred carbon source of *S. cerevisiae* [48]. Such mutational robustness of promoter activity could be the result of adaptation to a commonly experienced environment [60-63] or fluctuating selection [64, 65], although nonadaptive processes could also account for this result [66-69]. Because mutations create the phenotypic variation necessary for the action of other evolutionary forces and because organisms are constantly faced with environmental changes, differences in the distributions of mutational effects among environments can play an important role in trait evolution [12, 70-72].

Focusing on the effects of individual mutations, we found that approximately 10-30% of the 235 mutations examined showed significant GxE affecting *P_TDH3_* activity, depending on the pair of environments. This observation indicates that mutations in the *TDH3* promoter readily generate genetic variation affecting expression plasticity, providing the raw material needed for natural selection to alter plasticity [7, 73, 74]. The mutations tested in *P_TDH3_* increased the difference in expression between glucose and galactose more than the polymorphisms examined, suggesting that natural selection has eliminated alleles that increase the plasticity beyond a certain degree.

Because only two mutations significantly reduced plasticity between this pair of environments, we were unable to determine whether reduced plasticity between glucose and galactose is more likely to be neutral, deleterious or beneficial. Therefore, we cannot distinguish between a hypothesis of directional selection favoring minimal plasticity and a hypothesis of stabilizing selection maintaining a particular degree of plasticity. The wild type allele and all but one of the 30 polymorphisms examined were recently shown to confer maximal fitness in the glucose-based medium tested [50], but the fitness effects of *P_TDH3_* alleles will also need to be determined in galactose to discriminate between these two different evolutionary scenarios.

In addition to advancing our understanding of how mutation and selection interact to maintain expression plasticity in natural populations, the single nucleotide resolution of our data allowed us to identify potential molecular mechanisms underlying plasticity and gene-by-environment interactions of the *TDH3* promoter. These data suggest that the condition-dependent use of transcription factor binding sites is primarily responsible for differences in regulation of *TDH3* expression between fermentable and non-fermentable environments whereas differences in chromatin structure might play a larger role in generating plasticity between the two fermentable environments tested. Prior studies hypothesizing mechanisms for gene-by-environment interactions have focused primarily on variation in context-dependent *cis*-regulatory sequences [19, 21, 75]. Characterizing the distributions of mutational effects in multiple environments for other promoters, including mutations in the *trans*-acting factors that interact with these promoters, will be necessary to draw general conclusions about the mechanisms affecting the evolution of gene expression plasticity.

## Materials and Methods

### Yeast strains

Construction of the un-mutagenized reference strain YPW1 containing the *P_TDH3_*-*YFP* reporter gene is described fully in Gruber et al. [49]. This reporter gene contains a 678bp *TDH3* promoter (*P_TDH3_*) sequence fused to the coding sequence for YFP Venus optimized for expression in *S. cerevisiae* [76] and the *CYC1* (cytochrome c isoform 1) terminator and is inserted near the *SWH1* pseudogene on chromosome 1 of strain BY4724 [77] at position 199270. All strains included in this study were derived from BY4724 and therefore are auxotrophs for uracil and lysine and haploids with *MAT**a*** mating type. Construction of 236 mutant strains, each containing a single G:C→A:T transition in the *TDH3* promoter region of this reporter gene, was accomplished by PCR-mediated site-directed mutagenesis, as described in Metzger et al. [42]. One of these strains, YPW519, was excluded from this study because it appeared to have acquired a secondary mutation that increased expression of *P_TDH3_*-*YFP.* To identify natural haplotypes of the *TDH3* promoter, the 678 bp promoter region was sequenced in 85 isolates of *S. cerevisiae* [78], identifying 28 distinct haplotypes [42]. These haplotypes differed from each other by one to 13 polymorphisms. Each of these haplotypes was PCR-amplified and inserted upstream of the *YFP* coding sequence in the YPW1 genetic background, as described in Metzger et al. [42]. 15 additional haplotypes of *P_TDH3_* that differed from one of the 28 natural haplotypes by only a single polymorphism were then constructed by PCR-mediated site directed mutagenesis of a natural haplotype and cloned upstream of the *YFP* coding sequence. Together, these 15 haplotypes plus the 28 haplotypes observed among the isolates sampled resulted in a set of 39 *P_TDH3_* haplotypes in which each haplotype differed from at least one other haplotype by as little as one polymorphism. The remaining 4 haplotypes, all of which were observed in one or more natural isolates, differed from the most similar haplotype by more than one polymorphism and were excluded from this study. Relationships among these haplotypes were represented in a haplotype network as described below and in Metzger et al. [42]. In all, these 39 haplotypes contained 30 unique polymorphisms in the *TDH3* promoter (Dataset S2).

### Growth conditions

The fluorescence levels of strains included in this study were quantified in four consecutive experiments. The goal of the first experiment was to quantify the plasticity of expression of the wild type *P_TDH3_* allele by measuring (in parallel) fluorescence of strain YPW1 after growth in media containing glucose, galactose or glycerol as carbon source. The other three experiments measured fluorescence of the 236 *cis*-regulatory alleles with point mutations in *P_TDH3_* and the 39 *P_TDH3_* haplotypes with different polymorphisms in each of the three carbon source environments. Fluorescence data for samples grown in rich media containing glucose originally collected for and described in Metzger et al. [42] were re-analyzed for this study using scripts modified to allow identical analyses in multiple environments.

We started the first experiment by reviving strain YPW1 that contained the wild type *P_TDH3_*-*YFP* reporter gene and strain BY4724 used to correct for autofluorescence from frozen glycerol stocks onto YPG agar plates (10 g/l Yeast extract, 20 g/l peptone, 3% v/v glycerol and 20 g/l agar). After 48 hours of growth at 30°C, we filled 12 wells of a deep 96-well plate with 0.5 ml of liquid YPG, 12 wells with 0.5 ml of YPD (10 g/l yeast extract, 20 g/l peptone and 20 g/l D-glucose) and 12 wells with 0.5 ml of YPGal (10 g/l yeast extract, 20 g/l peptone and 20 g/l galactose). Other treatments were applied to the remaining wells, but these data were not used in this study (see “Layout.Plasticity.txt” worksheet in Dataset S3 for full description). Six wells containing YPG were inoculated with strain YPW1 and the six other wells were inoculated with strain BY4724 to a density of ~1.5 × 10^−5^ cells/ml. The plate was then incubated at 30°C with constant orbital shaking at 250 rpm. Each well contained a 3 mm glass bead that maintained cells in suspension during growth. After 22 hours of growth, YPW1 and BY4724 were each inoculated in six wells containing YPD and six wells containing YPGal to a density of ~1.5 × 10^5^ cells/ml. The plate was then incubated for another 20 hours at 30°C with constant shaking. We delayed the inoculation of samples in YPD and YPGal so that all samples reached a density above 6 × 10^7^ cells/ml simultaneously a few hours before the end of the experiment, despite the slower growth rate in the non-fermentable YPG (~170 min/division) as compared to the fermentable YPD (~85 min/division) and YPGal (~95 min/division). Samples were grown in parallel in the three carbon source environments to avoid previously observed day-to-day variation in flow cytometer sensitivity that complicates quantitative comparisons of absolute fluorescence levels between experiments. After growth, samples were diluted to ~2.5 × 10^6^ cells/ml in a clean plate containing 0.5 ml per well of synthetic complete medium lacking arginine (with the same carbon source used for growth) and fluorescence levels of single cells were quantified by flow cytometry as described below.

For the three experiments performed separately in media containing glucose, glycerol or galactose, yeast strains were arrayed and kept frozen at −80°C in eight 96-well plates prior to use. These experiments included the 236 mutants strains with point mutations in the *TDH3* promoter driving expression of YFP, the 39 strains with natural and reconstructed haplotypes of the *TDH3* promoter driving expression of YFP (used to quantify the effects of 30 unique polymorphisms on fluorescence levels), as well as appropriate controls such as YPW1 with the wild type allele of *P_TDH3_* driving expression of YFP and BY4724 used to correct for autofluorescence. Other strains not used for this study were also included in these experiments. Importantly, each plate contained 20 replicates of the control strain YPW1 at fixed positions used to correct for technical variation in fluorescence levels. Apart from these controls, the position of other strains was fully randomized to avoid systematic positional bias in fluorescence levels for different types of mutants (see “Layout.Mutations.Polymorphisms” worksheet in Dataset S3 for full description). Prior to each experiment, samples from each of the eight frozen plates were transferred to an OmniTray containing YPG agar medium. After 48 hours of growth, samples from each OmniTray were transferred with a V&P Scientific pintool to three 96-well plates containing either 0.5 ml of YPD (glucose), 0.5 ml of YPG (glycerol) or 0.5 ml of YPGal (galactose) per well (with one 3 mm glass bead per well), for a total of 24 plates in each experiment (three replicates of the eight initial plates). Plates were incubated at 30°C with constant shaking at 250 rpm during 20 hours for samples grown in YPD or YPGal and during 42 hours for samples grown in YPG. After growth, 20 μl of cell culture was diluted into 0.5 ml of synthetic complete medium lacking arginine (with the same carbon source used for population growth) in a clean 96-well plate that was immediately run through the flow cytometer for quantification of fluorescence levels as described below. 24 additional 96-well plates were inoculated the next day from the same OmniTrays stored at 4°C. These samples were grown and their fluorescence scored following the same procedure as described above, so that in total fluorescence was measured for six replicates of the eight initial plates (*i.e.*, six replicate populations of each mutant strain). For the experiment performed in YPD, we quantified fluorescence for nine replicate populations of each sample, but only analyzed data from six replicates to keep the number of replicates consistent between environments. For the experiment performed in YPGal, two plates were accidently flipped during manipulations required for the experiment and data associated with these plates was excluded from our analysis.

### *Quantification of* cis-*regulatory activity*

Activity of the *P_TDH3_*-*YFP* reporter gene was measured by flow cytometry using similar experimental procedures as in Metzger et al. [42] and a similar analysis pipeline as in Duveau et al. [50]. After dilution in SC-Arginine, cells were directly sampled from 96-well plates using an IntelliCyt HyperCyt Autosampler and passed through a BD Accuri C6 flow cytometer using a flow rate of 14 μl/min and core size of 10 μm. A blue laser (λ = 488 nm) was used for excitation of YFP and fluorescence was acquired from the FL1 channel using a 533/30 nm optical filter. Liquid cultures of each strain were sampled for 2-3 s each, with ~20,000 events recorded on average. Data was then processed with custom scripts using *flowClust* (3.0.0) and *flowCore* (1.26.3) packages in R (3.0.2) to remove artifacts such as debris and other non-cell events [79, 80] as well as cell doublets. Samples with fewer than 800 events following processing were excluded from further analysis. Next, we defined a measure of fluorescence level that was independent of cell size, which was complicated by the fact that the positive relationship between fluorescence intensity (FL1.A) and cell size (forward scatter, FSC.A) was not the same for samples collected in glucose, glycerol and galactose. To correct for cell size homogeneously in the three environments, we transformed the log_10_(FSC.A) and log_10_(FL1.A) data for each sample using a rotation around the centroid and with an angle determined iteratively so that the intercept of the linear regression of the transformed values of log_10_(FSC.A) and log_10_(FL1.A) would be close to zero (below 0.01). The fluorescence level for each event was then calculated as the ratio of the transformed value of log_10_(FL1.A) over the transformed value of log_10_(FSC.A) and the fluorescence level for each sample was calculated as the median fluorescence across all events (or cells) in the sample.

For the experiment designed to determine the plasticity of activity of the wild type promoter, we subtracted the average autofluorescence measured for the six replicates of non-fluorescent strain BY4724 in each environment from the fluorescence levels measured for strain YPW1 (wild type *P_TDH3_*-*YFP*) in the corresponding environments. After autofluorescence correction, we divided the median fluorescence measured for each YPW1 replicate in each environment by the average fluorescence measured for the six replicates of YPW1 in YPD (glucose). We finally calculated the mean relative fluorescence across the six replicates of YPW1 in each environment to determine the plasticity of activity of the wild type *TDH3* promoter. The custom script describing the plasticity analysis is provided as Dataset S4.

For the other experiments that included a larger number of plates, control samples were used to test and correct for technical variation creating differences among days and plates as well as rows and columns within a plate. To do this, we fit YFP fluorescence from the control samples (median log_10_(FL1.A)/log_10_(FSC.A) across all cells) to a linear model (fluorescence ~ day + run + replicate + plate + row + column + block + stack + depth + order) using the *lm* function in R. Effects in this model correspond to the day in which a plate was run, the replicate number for that plate in that day, the run that uniquely identified each physical plate run on the flow cytometer, the plate number corresponding to one of the eight arrays, row, column, and block in which a sample was run (due to software limitation, data was acquired from each plate in multiple blocks), the stack in which a plate was cultured, its depth in this stack, and the order in which a plate was run within the replicate. The function *step* in R was used to choose the model with the best explanatory power based on AIC statistic and to drop the factors that did not significantly impact fluorescence. “run” and “block” were the main factors affecting fluorescence, therefore we used the linear model “fluorescence ~ day + run” to estimate the effects of these two factors and to correct the fluorescence levels of all samples accordingly. We then subtracted the average autofluorescence measured across the six replicates of nonfluorescent strain BY4724 from the fluorescence of all samples. Finally, the fluorescence of each sample was divided by the mean fluorescence measured across replicates of the reference strain YPW1 in the same environment. The mean relative fluorescence measured across the six replicates of each strain was used as a measure of the effects of the 235 mutations on *TDH3* promoter activity in glucose, galactose and glycerol. Importantly, our measure of mutational effects was independent of the plasticity of expression observed between environments, allowing us to test for gene-by-environment interactions simply by comparing the effects of a mutation between two environments. The custom script used to perform these analyses is provided as Dataset S5.

### Effects of individual polymorphisms

The effects of polymorphisms were measured in each environment as described in Metzger et al. [42] and Duveau et al. [50]. Using parsimony, a haplotype network (see “Haplotype.Network.txt” worksheet in Dataset S3) was generated for the 39 *P_TDH3_* haplotypes described above that each differed from another haplotype by exactly one polymorphism. The most likely ancestral state of the *TDH3* promoter was then inferred using the *TDH3* promoter sequences of other species in the *Saccharomyces senso strictus* genus and more distantly related strains of *S. cerevisiae* and used to polarize the network [42]. Parsimony and maximum likelihood methods were used to construct this haplotype network. Conservation across the 678 bp *TDH3* promoter shown in Figure 4 and Figure S1 was determined by comparing sequences of the seven species of the *Saccharomyces stricto sensus* genus in 20 bp sliding windows using ConSurf [42, 81, 82].The activity of these promoter haplotypes was measured in parallel with the activity of the 235 mutant promoters after growth in YPD, YPG and YPGal as described above. After correcting for autofluorescence, we divided the fluorescence measured for each replicate of each haplotype by the average fluorescence across the six replicates of the parental haplotype that differed by a single polymorphism in the haplotype network. The mean relative fluorescence across the six replicates of each haplotype represented the effects of individual polymorphisms on promoter activity. In seven instances, the effect of the same polymorphism was tested in two different pairs of haplotypes. In such cases, after verifying the absence of epistatic interaction [50], we averaged the effects of the polymorphism measured with the two pairs of haplotypes. Overall, this approach allowed us to quantify the effects of 30 unique polymorphisms in the *TDH3* promoter. 10 of these polymorphisms were G:C to A:T transitions (the same type of change tested in the collection of 235 mutations), 13 were other types of single nucleotide polymorphisms and 7 were small indels ranging from 1 to 13 bp. We observed no significant difference of effects among these three classes of polymorphism in any environment (Figure S1B-D).

### Statistical analyses

All statistical tests were performed using R 3.2.3. To compare the activity of the wild type *TDH3* promoter in different environments, we performed t-tests on the fluorescence levels measured for the six replicate populations of YPW1 in each pair of environments. To compare the distribution of mutational effects for different pairs of environments, we performed Kolmogorov-Smirnov tests using R function *ks.test* followed by 200,000 permutations to determine the distribution of D statistics (supremum difference between the two empirical distribution functions) expected under the null hypothesis of no plasticity. For each permutation, we shuffled the two environment labels in which the effects of each of the 235 mutations were measured (the number of possible permutations was 2^235^) and the D statistic was calculated for the two randomized distributions. The P-value of the KS test was determined as the proportion of the 100,000 permuted distributions that showed a greater D statistic than the one obtained from the observed distributions of mutational effects.

We used non-parametric permutation tests to compare the median effects of mutations for different pairs of environments. For each permutation, we generated two random sets of 235 mutational effects by shuffling the effects measured in the two environments of interest for each mutation (the number of possible permutations was 2^235^). We then calculated the difference of median effects between the two random sets and repeated the procedure 200,000 times to generate a null distribution of differences of medians. The two-sided P-value of the test was calculated as twice the proportion of random sets that showed a more extreme difference of median effects than the observed difference between the two environments. We used similar permutation tests to compare the median absolute deviation (MAD; a measure of dispersion) of mutation effects between environments. To do so, for each pair of environments, we first centered the effects of the 235 mutations on the same median by subtracting the median effects of all mutations observed in one environment to the effect of each mutation measured in the other environment. This transformation did not affect the variability of mutational effects (MAD) and ensured that the permutation test was unaffected by differences in median effects of mutations between environments. We then used the same permutation procedure as described above to generate 200,000 differences of MAD between randomized sets of mutation effects and calculated P-values as twice the proportion of random MAD differences that were more extreme than the observed difference of MAD across all 235 mutations between the two environments.

We tested for significant gene-by-environment interactions using t-tests (*t.test* function in R) to compare the effect of each mutation in each pair of environments. A Benjamini-Hochberg false discovery rate correction was then applied to control for multiple testing, and mutations with adjusted P-values below 0.01 were considered to show statistically significant GxE effects (*i.e.*, these mutations showed different effects in the two environments tested). We compared the magnitude of GxE effects across the 235 mutations between pairs of environments using Mann-Whitney-Wilcoxon tests (*wilcox.test* function in R).

We tested for evidence of selection acting on the average activity of the *TDH3* promoter in each environment or on the plasticity of activity between environments using the method developed in Metzger et al. [42]. First, the distributions of effects of mutations (or the distributions of differences of effects of mutations between environments) were converted into probability density functions. Then, these functions were used to compute the log-likelihood of 200,000 sets of 30 mutational effects randomly drawn with replacement from the empirical distributions of 235 mutations effects. The log-likelihood value for the effects (differences of effects between environments) of the 30 polymorphisms was then calculated and a two-sided P-value was calculated as twice the proportion of random sets of mutational effects showing a more extreme log-likelihood value than the one observed for polymorphisms. P-values below 0.05 indicated that the effects of the 30 polymorphisms were statistically different from the effects of a random set of 30 mutations, suggesting a role of natural selection in shaping the phenotypic effects of polymorphisms. In addition to this test for selection, we also compared the median effects of polymorphisms to the median effects of random sets of 30 mutations using permutation tests. For these tests, we randomly picked a set of 30 mutational effects (or 30 differences of mutational effects between environments) and calculated the difference *A* between the median effect for this random set and the median effect measured across the 30 polymorphisms. The effects of the 30 mutations and 30 polymorphisms were then shuffled and a randomized difference *B* of median effects between the two permuted sets was calculated. Lastly, we calculated the difference *D* equal to *A* – *B.* After 200,000 repetitions of this procedure, we calculated the frequency of *D* values that were above zero and the frequency of *D* values that were below zero. The two-sided P-value of the test was calculated as twice the minimal frequency.

A nearest neighbor approach was used to test for non-random positioning of GxE effects in the *TDH3* promoter. First, we determined the positions in the promoter of mutations with significant differences of effects between two environments. For each of these mutations, we calculated the distance to the closest mutation with significant GxE effect in the promoter (nearest neighbor). Then, we computed the standard deviation of the minimal distance across all mutations with significant GxE effects between the two environments. Lastly, similar standard deviations of minimal distance were calculated for 100,000 random sets of mutations (picked among the 235 mutations created in the promoter). The number of random mutations drawn for each set was equal to the number of mutations with significant GxE effects for which non-random positioning was being tested. Two-sided P-value of the test was calculated as twice the proportion of randomized standard deviations of distance to nearest neighbor that were more extreme than the observed standard deviation. P-value below 0.05 can be obtained either if the mutations with GxE effects tend to cluster in the promoter or if they tend to be more homogeneously spaced than expected by chance.

We used a resampling method to test for enrichment or depletion of GxE effects in functional elements of the *TDH3* promoter. 100,000 sets of mutations of the same size as the number of significant GxE effects were randomly drawn without replacement from the mutations we created in the promoter, excluding mutations in functional elements that were not being tested. We then calculated the proportion P of random sets of mutations that contained more mutations in the tested functional element than the observed number of GxE effects falling in this functional element. The P-value of the test was calculated as twice the lowest value between P and 1-P.

### Access to data and analysis scripts

Flow cytometry data used in this work is available through the Flow Repository (https://flowrepository.org). Repository ID = FR-FCM-ZZBN contains data from Metzger et al. (2015) used to quantify the effects of the mutations and polymorphisms in *P_TDH3_* on expression after growth in YPD (glucose), repository ID = FR-FCM-ZY8D contains data collected to analyze expression of these genotypes in YPGal (galactose), repository ID = FR-FCM-ZY8B contains data collected to analyze expression of these genotypes in YPG (glycerol) and repository <pending> contains data used to quantify the plasticity of activity of the wild type *TDH3* promoter in glucose, galactose and glycerol. The R scripts used to perform the analyses described above are provided as Datasets 4 and 5. Data used by these scripts are provided in Dataset S3 and data produced by these scripts are included in Dataset S1.

## Acknowledgements

We thank Jonathan Gruber for technical assistance as well as Petra Vande Zande, Jennifer Lachowiec, Abigail Lamb and Rebecca Tarnopol for comments on the manuscript. Funding for this work was provided to P.J.W. by the March of Dimes (5-FY07-181), Alfred P. Sloan Research Foundation, National Science Foundation (MCB-1021398), National Institutes of Health (1R01GM108826, 1R35GM118073) and the University of Michigan. Additional support was provided by National Institutes of Health Genetics training grant (T32 GM007544) to D.C.Y.; the University of Michigan Rackham Graduate School, Ecology and Evolutionary Biology Department and the National Institutes of Health Genome Sciences training grant (T32 HG000040) to B.P.H.M.; EMBO postdoctoral fellowship (EMBO ALTF 1114-2012) to F.D.; and National Institutes of Health National Research Service Award (1F32GM115198) to A.H-D.

---

**Figure S1.**
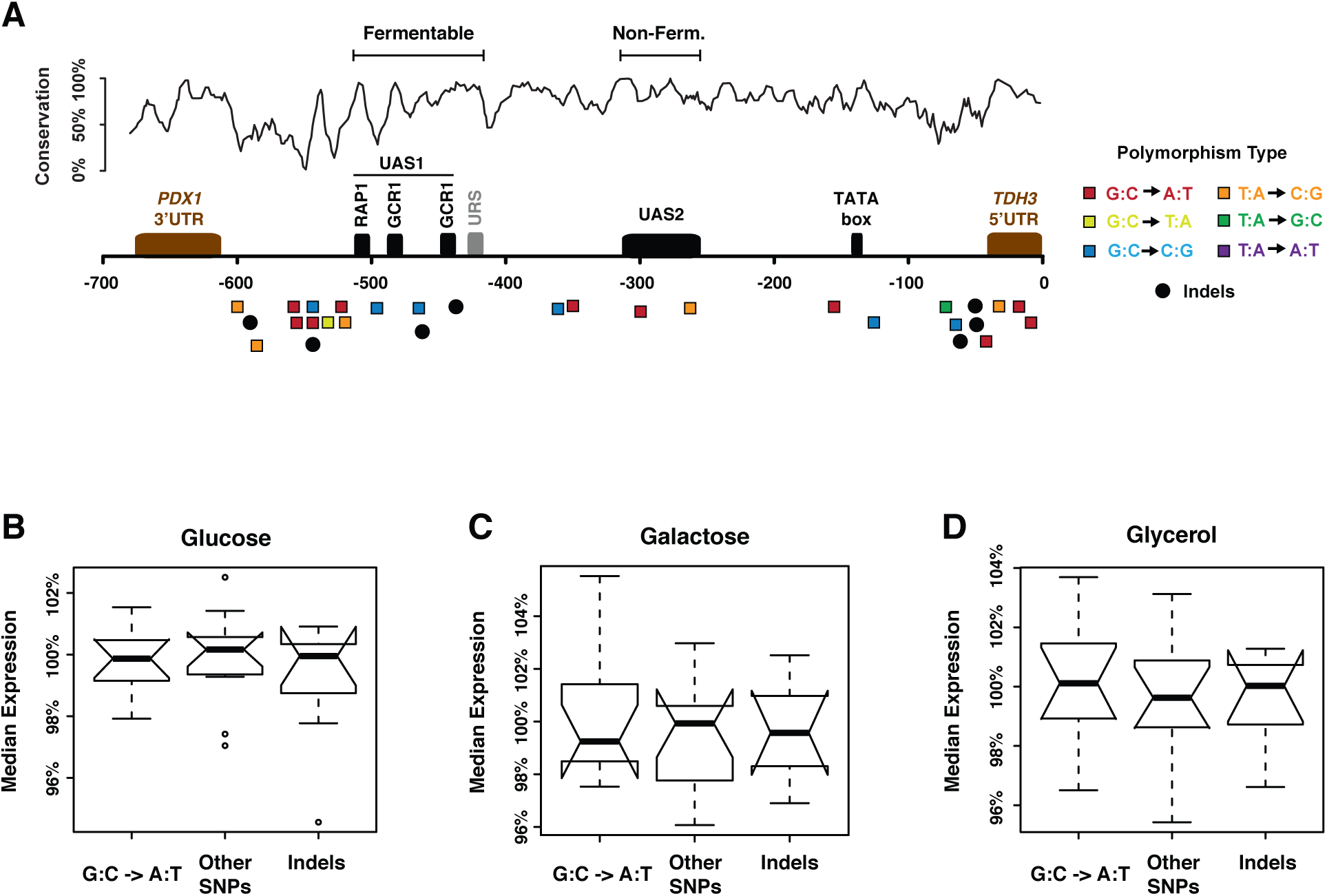
Polymorphisms in the *TDH3* promoter. (A) Schematic shows the positions and types of the 30 unique polymorphisms examined in this study. Single nucleotide polymorphisms (SNPs) are indicated with squares of different colors corresponding to the type of base change (see legend on the right). Small indels are indicated with black circles. Key functional elements are also shown, with UAS1, UAS2, and URS discussed in the main text. Note that none of the polymorphisms affected the RAP1 or GCR1 binding sites or the TATA box. The black curve shows sequence conservation across species of the *Saccharomyces senso stricto* genus. (B-C) Boxplots comparing the effects of different types of polymorphisms on *P_TDH3_* activity upon growth in (B) glucose, (C) galactose, or (D) glycerol are shown. Each box includes the average effects of 6 replicates for the 10 G:C to A:T transitions, 13 other kind of SNPs or 7 indels shown in (A). Thick horizontal lines represent median effects across all polymorphisms of each type in each environment. Notches indicate the 95% confidence interval of the medians and boxes show interquartile range. Mann-Whitney-Wilcoxon tests were performed to compare the average effects of different types of polymorphisms and resulted in non-significant P-values (P > 0.5) for all pairwise comparisons.

**Figure S2.**
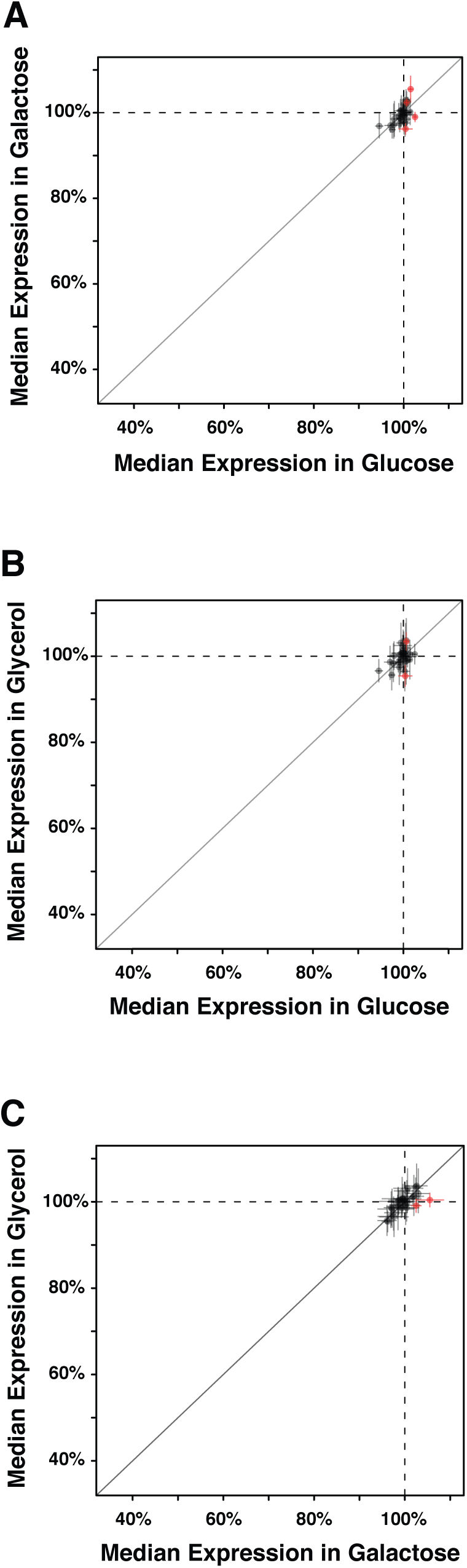
Pairwise comparisons of the effects of polymorphisms on *P_TDH3_* activity in different environments. (A-C) The median effect of each polymorphism on *P_TDH3_* activity relative to the ancestral haplotype used to infer its effect is compared between (A) glucose and galactose, (B) glucose and glycerol and (C) galactose and glycerol. Dots indicate the effect of polymorphisms in each environment calculated as the ratio of the mean fluorescence (across six replicates) of pairs of haplotypes that differed by a single polymorphism. Error bars show 95% confidence intervals calculated using Fieller’s theorem. Red dots indicate polymorphisms with no overlap of the 95% confidence intervals between the two environments considered, which were interpreted as showing significant gene-by-environment interactions.

**Figure S3.**
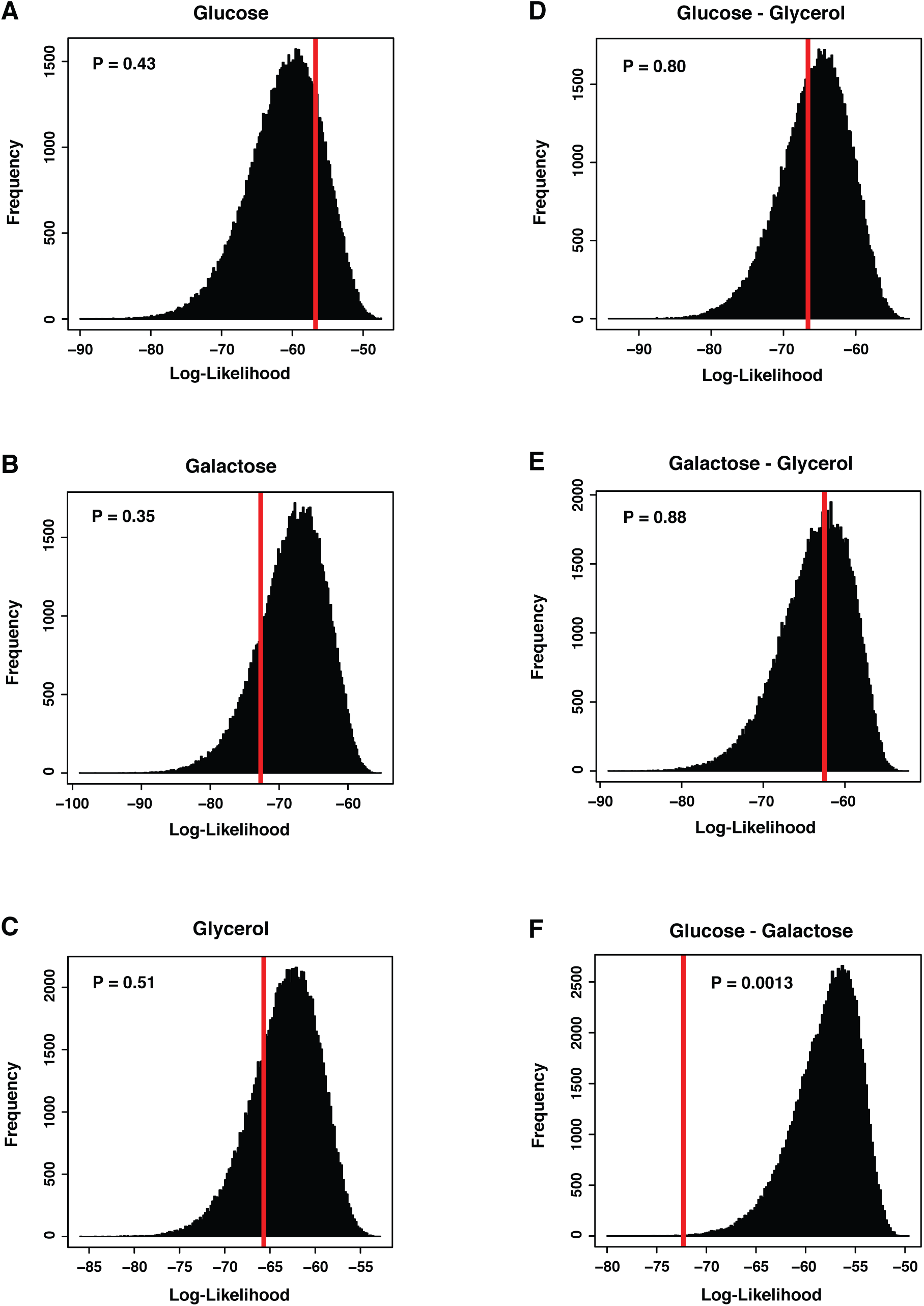
Testing for evidence of selection acting on *PTDH3* activity and plasticity among environments. (A-C) Histograms showing the distribution of log-likelihood for 100,000 random sets of 30 mutational effects drawn with replacement from the 235 effects of mutations measured in (A) glucose, (B) galactose and (C) glycerol. Two-sided P-values represent twice the proportion of random sets of 30 mutation effects with a log-likelihood more extreme than the log-likelihood value calculated from the effects of the 30 polymorphisms in each environment (red lines). A low P-value indicates that the distribution of effects of polymorphism is different from random sampling of mutational effects. (D-F) Histograms showing the distribution of log-likelihood for 100,000 random sets of 30 GxE effects randomly drawn with replacement from the 235 GxE effects measured between (A) glucose and glycerol, (B) galactose and glycerol, and (C) glucose and galactose. The red lines indicate the log-likelihood value calculated from the GxE effects measured for the 30 polymorphisms between each pair of environments. Two-sided P-values were calculated as twice the proportion of random GxE effects with more extreme log-likelihood than the log-likelihood calculated from the GxE effects of polymorphisms. Low P-values indicate that the plasticity of expression caused by polymorphisms differs from the plasticity sampled from random mutations.

**Dataset S1. Output data files produced by analysis**

**Dataset S2. Summary of naturally occurring *P_TDH3_* haplotypes and polymorphisms**

**Dataset S3. Input data files used for analysis**

**Dataset S4. R script used to analyze flow cytometry data testing for plasticity of the wild type *PTDH3* allele**

**Dataset S5. R script used to analyze flow cytometry data estimating and comparing the effects of mutations and polymorphisms on *P_TDH3_* activity**

